# Structure of *Aquifex aeolicus* Lumazine Synthase by Cryo-Electron Microscopy to 1.42Å Resolution

**DOI:** 10.1101/2024.03.21.586070

**Authors:** Christos G. Savva, Mohamed A. Sobhy, Alfredo De Biasio, Samir M. Hamdan

**Affiliations:** Division of Biological and Environmental Sciences and Engineering, King Abdullah University of Science and Technology, Thuwal, Saudi Arabia

## Abstract

Single particle Cryo-Electron microscopy (Cryo-EM) has become an essential structural determination technique with recent hardware developments making it possible to reach atomic resolution at which individual atoms, including hydrogen atoms, can be resolved. Thus Cryo-EM allows not only unprecedented detail regarding the structural architecture of complexes but also a better understanding surrounding their chemical states. In this study we used the enzyme involved in the penultimate step of riboflavin biosynthesis as a test specimen to benchmark a recently installed microscope and determine if other protein complexes could reach a resolution of 1.5Å or better which so far has only been achieved for the iron carrier ferritin. Using state of the art microscope and detector hardware as well as the latest software techniques to overcome microscope and sample limitations, a 1.42Å map of *Aquifex aeolicus* lumazine synthase (AaLS) was obtained from a 48-hour microscope session. In addition to water molecules and ligands involved in AaLS function, we can observe positive density for ∼50% of hydrogen atoms. A small improvement in resolution was achieved by Ewald sphere correction which was expected to limit the resolution to ∼1.5Å for a molecule of this diameter. Our study confirms that other protein complexes can be solved to near-atomic resolution. Future improvements in specimen preparation and protein complex stabilization may allow more flexible macromolecules to reach this level of resolution and should become a priority of study in the field.

## Introduction

Single particle Cryo-EM has advanced at a fast pace over the last 10 years beginning with the “Resolution Revolution” in 2013 (Bai *et al*., 2013). The exponential growth of the field is highlighted by the number of deposited structures in the Electron Microscopy Database (EMDB) which surpassed 30K cumulative maps in the year 2023 and with almost 25% of these being deposited during that same year (Source: EMDB statistics; March 2024). This highlights the adoption of Cryo-EM as a mainstream structural biology determination approach by existing and newly established structural biology groups around the world. Furthermore, the rapid development of user-friendly software for high-speed data collection and the creation of National or Regional Facilities worldwide have made the process of collecting high-quality data more tangible than ever before. Recently the potential of a low-cost screening/data collection microscope was demonstrated which will enable the dissemination of the technique to more researchers (McMullan *et al*., 2023).

While significant hurdles still exist in obtaining suitable samples for Cryo-EM with few cases being straight forward and requiring an optimization process by trial and error, the potential of atomic-resolution SPA (i.e. close to 1.2Å) was demonstrated on the gold-standard apoferritin by several groups beginning in 2020 (Nakane *et al*., 2020, Yip *et al*., 2020). The recombinant version of either mouse or human ferritin can routinely reach resolutions higher than 2Å on Field Emission Gun (FEG) microscopes while the use of a highly-coherent Cold Field Emission Gun (cFEG) or the use of a monochromator and spherical aberration corrector greatly increase the signal attainable near the 1Å resolution range (Nakane *et al*., 2020). The use of these latest hardware allowed the structures of apoferritin to be determined to 1.22Å and 1.25Å respectively while in 2023 a 1.19Å map was also reported (Maki-Yonekura *et al*., 2023). This leap in resolution resulted in maps in which individual atoms could be placed unambiguously and the visualization of hydrogen atom positions by calculating difference maps between the experimental density and atomic coordinates possible (Yamashita *et al*., 2021). More recently other (non-test specimen) SPA derived structures have also approached near-atomic resolution including the 1.55Å structure of a prokaryotic ribosome (Fromm *et al*., 2023) and the 1.52Å structure of *M. smegmatis* Huc complex (Grinter *et al*., 2023).

The Imaging and Characterization Core lab at the King Abdullah University of Science and Technology (KAUST) acquired a Thermo-Fisher Scientific (TFS) Krios G4 in the summer of 2022 to enable high-throughput Cryo-EM for the users in the University and wider region. During commissioning of the microscope, a 2.0Å map of mouse apoferritin was obtained using aberration free Image shift (AFIS). The Krios G4 equipped with a cFEG source, Selectris-X post-column energy filter and a Falcon 4i detector is almost identical to the setup which resulted in the 1.22Å apoferritin structure and therefore we wanted to repeat a benchmark to identify any potential issues with this microscope and explore its capabilities.

A previous study conducted locally on an older Krios G1 upgraded with a Gatan Imaging Filter and a Gatan K2 detector explored the suitability of two high-symmetry protein complexes as benchmark candidates (Sobhy *et al*., 2022). Of these, AaLS from the hyper-thermophilic bacterium *Aquifex aeolicus*, forms a 1MDa spherical capsid of 60 identical subunits with icosahedral symmetry (Zhang *et al*., 2001). Using this older hardware, a reconstruction of 2.3Å was achieved which was very close to Nyquist for the pixel size used. Therefore, we opted to use AaLS as a test specimen to evaluate the Krios G4.

## Results

Using the same preparation of AaLS used for the 2.3Å Krios G1 derived map, we prepared specimens using UltrAuFoil grids to minimize beam-induced particle movement (Russo & Passmore, 2014). Careful alignment of the microscope optics to minimize objective lens astigmatism and axial comma were performed on grids coated with holey carbon immediately prior to data collection setup. The choice of suitable squares is very important when trying to obtain the best performance out of the microscope and detector as increased ice thickness can be detrimental to high-resolution structural determination. Several studies have been conducted to optimize square and hole selection using different aspects that are affected by sample thickness such as loss of electrons by either inelastic scattering or high-angle scattering through the objective aperture. These measurements, combined with experimental determination of ice thickness, can be used to calibrate on-the-fly ice thickness determination parameters (Rice *et al*., 2018, Rheinberger *et al*., 2021). Another report used Plasmon range energy-loss electrons to aid hole selection especially on gold foils where the increased thickness of the foil makes determining the ice thickness more challenging (Hagen, 2022). In practice, the ice thickness parameter must be taken into consideration alongside particle distribution and stability. In TFS EPU v3.5, a per-hole histogram function which relays the gray-level distribution in individual holes allows one to set the minimum and maximum grey-level range from such holes to the entire dataset. The reported dose in EPU can then be used to find the thinnest possible ice (by comparing to vacuum) that gives a good distribution of particles. We have been routinely using this approach when using UltrAuFoil grids to aid in hole selection and this approach was implemented for this dataset. From the resulting micrographs, 95% displayed CTF estimated frequencies better than 5Å and reaching ∼2.5Å in the best cases (Fig.1a,b)

**Figure 1.**
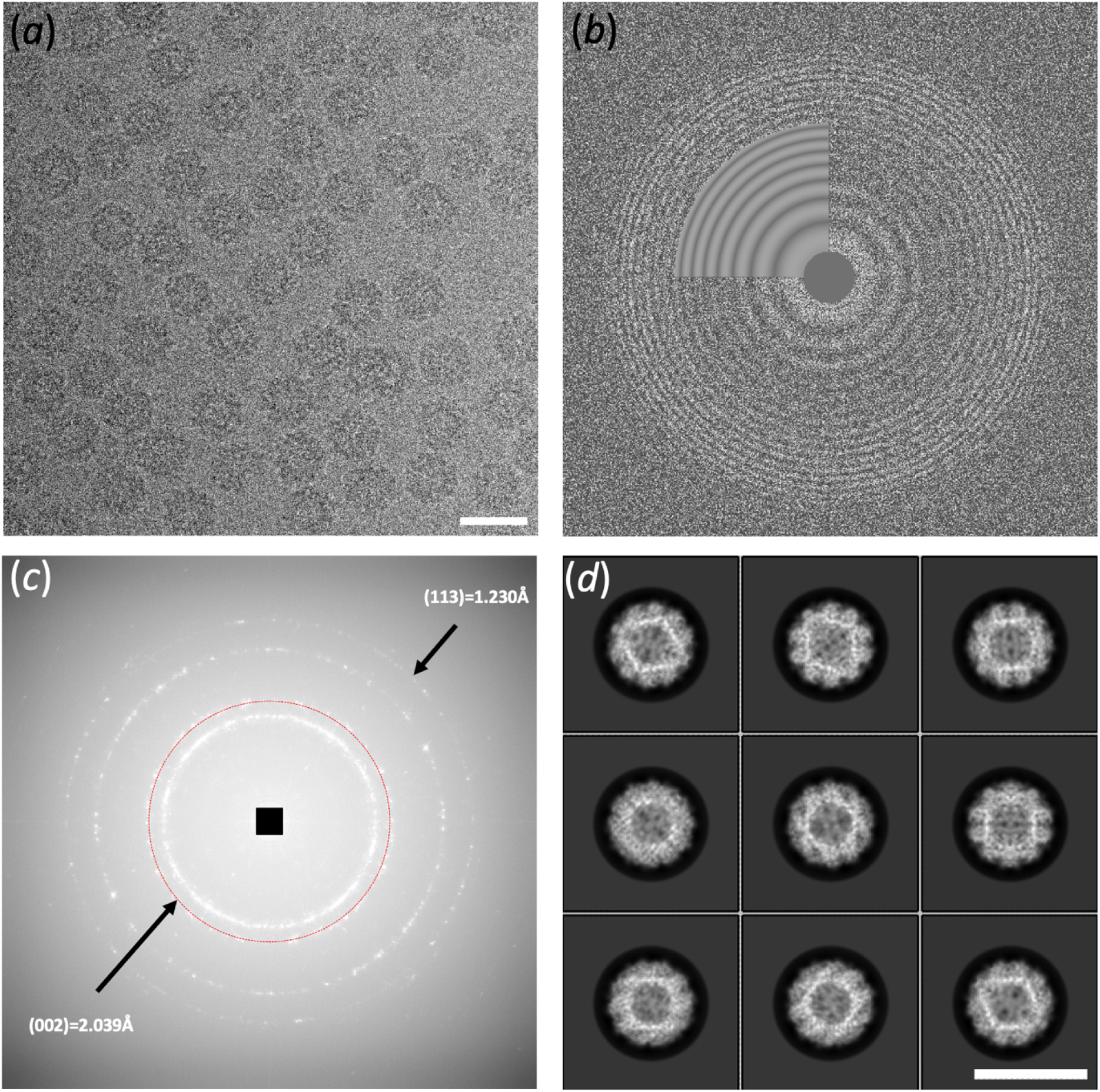
Cryo-EM of AaLS. (a) Reprsentative electron micrograph of AaLS particles taken at a defocus of ∼-1.0μm and a total dose of ∼46 e-/Å^2^. (b) Corresponding CTF parameter estimation power spectrum. (c) Summed power spectrum of 5 micrographs of UltrAuFoil support film indicating the (002) diffraction ring (red circle) used for pixel calibration. The (113) ring is also visible indicating frequencies to 1.23Å using conditions identical to data collection. (d) Class averages of 4x binned AaLS particles. Scale bars correspond to 200Å.

The calibrated pixel size is an important factor when reporting the resolution of cryo-EM data and especially when trying to obtain high resolution. In addition to causing fitting errors of the Contrast Transfer Function (CTF) at higher frequencies (Dickerson *et al*., 2024, Danev *et al*., 2021) the pixel size determines the reported resolution. Even if 4^th^ order aberration estimation is carried out during processing which can mitigate the effect of an incorrect pixel size (Zivanov *et al*., 2020), the input of the calibrated pixel size during post-processing is important for the correct scaling of maps to be used in model refinement and accurate reporting of resolution. As reported by others (Danev *et al*., 2021, Dickerson *et al*., 2024), the nominal calibrated pixel size which is determined by service engineers can be off by several percentiles. In addition, pixel calibration using grating replicas of varying ratios of metals such as gold/palladium, affects the position of the diffraction rings depending on this ratio and care should be taken to use pure metals for this purpose (Danev *et al*., 2021). Finally, the projection systems of microscopes are not tunable by the user and varying amounts of anisotropic magnification can be present from the factory (Zivanov *et al*., 2020, Grant & Grigorieff, 2015). The recently reported magCalEM software package has been developed specially for calibrating the correct pixel while taking into account the effect of anisotropic magnification (Dickerson *et al*., 2024). We used this software to obtain the correct pixel size which in our case was 1.2% larger than the service determined pixel size Fig.1c. The calculated power spectra of the UltrAuFoil support film indicated frequencies at least to 1.23Å (Fig.1c) under identical optics and dose conditions used for data collection.

We opted to use stage shift for this experiment to reduce the effect of comma on the datasets as reported before (Nakane *et al*., 2020). A square pattern of 9 exposures for each hole was used using a 350nm parallel beam with no contact between the beam and foil and no overlap of the illuminated areas. The dose-rate on the Falcon4i detector was set to ∼3 e-/pix/sec or approximately 1 electron per hundred pixels per frame to minimize coincidence loss (Greg McMullan; personal communication, TFS Falcon4i Applications Notes). Over the 48-hour session 12657 movies were recorded in EER format (Guo *et al*., 2020). Data collection parameters are summarized in Table 1.

**Table 1.**
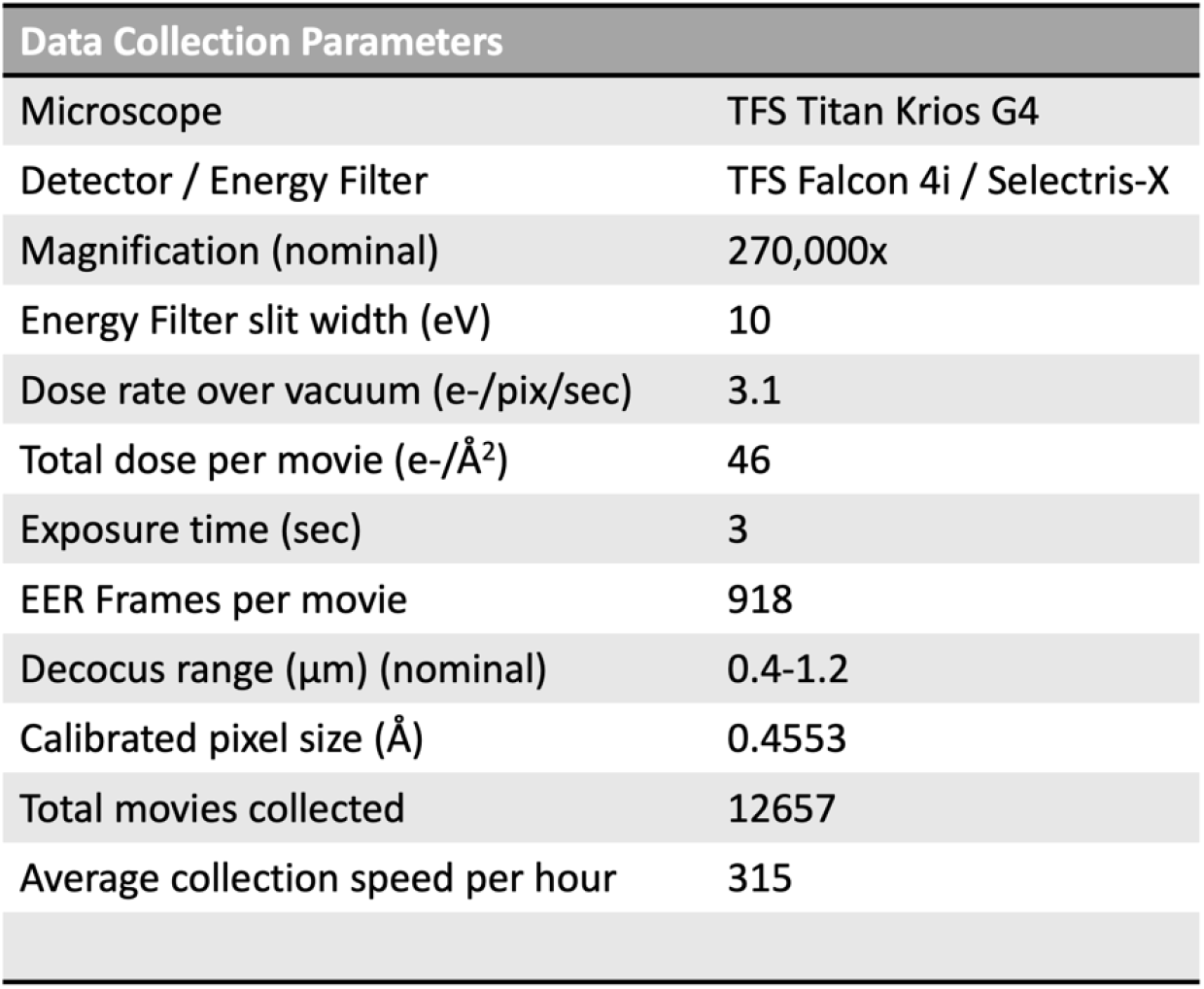
Cryo-EM data collection parameters.

| Data Collection Parameters |  |
| --- | --- |
| Microscope | TFS Titan Krios G4 |
| Detector / Energy Filter | TFS Falcon 4i / Selectris-X |
| Magnification (nominal) | 270,000x |
| Energy Filter slit width (eV) | 10 |
| Dose rate over vacuum (e-/pix/sec) | 3.1 |
| Total dose per movie (e-/Å<sup>2</sup>) | 46 |
| Exposure time (sec) | 3 |
| EER Frames per movie | 918 |
| Decocus range (μm) (nominal) | 0.4-1.2 |
| Calibrated pixel size (Å) | 0.4553 |
| Total movies collected | 12657 |
| Average collection speed per hour | 315 |

The final 3D reconstruction of AaLS consisted of 470,878 particles. This 1.43Å map was slightly improved by Ewald Sphere correction to 1.42Å (Fig2.a) (Sup. Fig. 1a,b) (Zivanov *et al*., 2018, Russo & Henderson, 2018). AaLS has a diameter of 160Å and the effect of the Ewald Sphere is expected to start affecting frequencies higher than 1.5Å (DeRosier, 2000). In this case it seems that the effect was not as prominent but it is possible that further improvement could be limited by the flexibility of the protein itself or the signal in the data. The Rosenthal B-Factor (Rosenthal & Henderson, 2003) was calculated to be 49Å^2^ using subsets of particles (Sup. Fig. 1d). The smallest subset consisting of 936 particles resulted in a 2.01 Å map.

**Figure 2.**
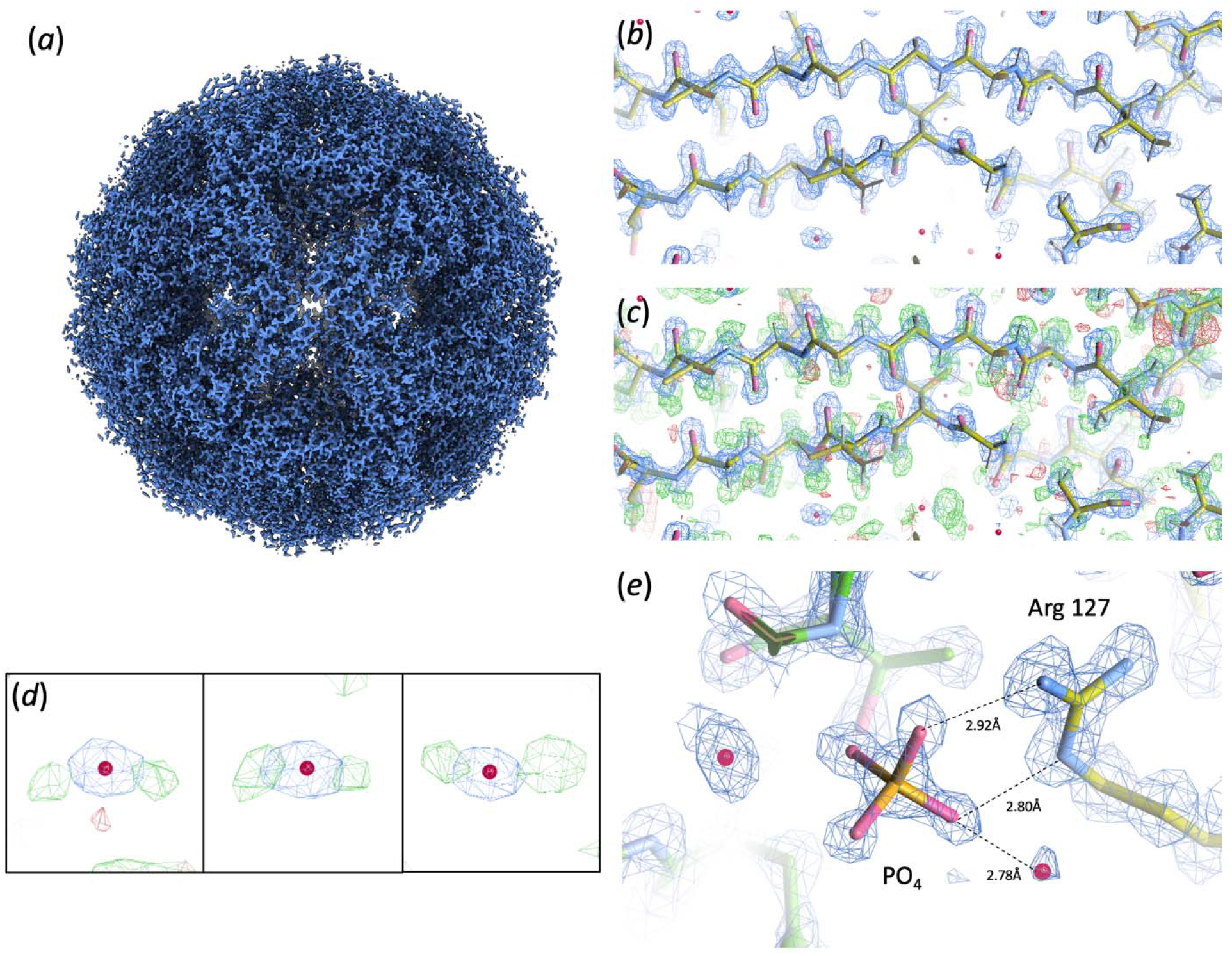
Map and model of AaLS. (a) Sharpened map of AaLS at 1.42Å resolution. (b) Density at 4.0s of a masked-normalised Fo map output from Servalcat. (c) Overlay of the Calculated *F*_o_*-F*_c_ difference map at 4.0s (b) indicating positive density (green) and in some areas negative density (red). Hydrogen atoms were built into the model if the corresponfing positive peaks were within 0.5Å of the putative H atoms. (d) Examples of identified water molecules with accompanying H atoms. (e) Arginine 127 interacting with a phosphate ion which is a bi-product of lumazine synthesis. Potential hydrogen-bonding distances indicated in dashed lines.

The half maps from Ewald Sphere correction were subsequently used for atomic model refinement using the 1.60Å crystal structure of AaLS as a starting model. Water molecules were identified in the sharpened map and of these, 43 were within 0.2Å of the water molecules modelled in the crystal structure (PDB ID: 1hqk). A mask-normalized *F*_o_*-F*_c_ generated difference map was then used to identify peaks within 0.5Å of hydrogen atoms in a reduced model, followed by manual curation which resulted in 552 hydrogens. Thus, we were able to identify ∼46% of all putative protein Hydrogen atoms from the map of AaLS at this resolution (Fig.2b,c). This number is consistent with previous studies using apoferritin maps determined at different resolutions. About 70% of hydrogen atoms could be located in maps at 1.19Å and 1.25Å resolution (Maki-Yonekura *et al*., 2023, Yamashita *et al*., 2021) while only ∼17% at 1.84Å resolution (Yamashita *et al*., 2021). Several of the water molecules identified displayed hydrogen densities adjacent to the central oxygen atom (Fig.2d). Finally, we observed density for a tetrahedral shaped ligand. This was built as a phosphate ion and is consistent as a bi-product of lumazine synthesis and in proximity to Arginine 127, a highly conserved residue in this family of proteins (Fig.2e)(Zhang *et al*., 2001). At pH 7.5 the phosphate ion is expected to be mostly in the HPO_4_^-2^ protonation state though we did not observe positive density for any hydrogens. It is unknown whether the phosphate is a result of enzymatic catalysis or simply carried over from phosphate present in the lysis buffer. Fourier shell correlation between the final map and model as well as cross-validation FSCs indicated no overfitting during refinement (Sup. Fig. 1c)(Brown *et al*., 2015). Model refinement statistics are shown in Table 2.

**Table 2.** Model refinement statistics.

| Model Refinement Statistics |  |
| --- | --- |
| Model resolution (Å) | 1.4 |
| Model resolution range (Å) | 1.4-180.30 |
| Mean overall B value (Å<sup>2</sup>) | 20.046 |
| Model composition |  |
| Non-hydrogen atoms | 1221 |
| Hydrogen atoms | 552 |
| Protein residues | 153 |
| Ligands | 1 |
| R.m.s. deviations |  |
| Bond lengths (Å) | 0.006 |
| Bond angles (°) | 1.456 |
| Validation |  |
| Clashscore (#/1000 atoms) | 1.7 (99<sup>th</sup> percentile) |
| Poor rotamers (%) | 0.85 |
| MolProbity score | 0.92 (100<sup>th</sup> percentile) |
| Ramachandran plot |  |
| Favored (%) | 98 |
| Allowed (%) | 2 |
| Outliers (%) | 0 |
| Cβ deviations (>0.25Å) | 0 |
| Cis Prolines | 0 |

## Discussion

In this study we set out to benchmark a recently installed, state of the art microscope and to identify any potential issues. The use of AaLS as a test sample allowed us to further examine its suitability as a viable benchmark specimen for higher-resolution structural determination. The structure we obtained from a 48-hour session at 1.42Å is to date the only sub-1.5Å structure of a complex other than apoferritin. Though apoferritin has long been used an ideal benchmark sample due to its stability, homogeneity and symmetry there is no reason why other complexes with the same characteristics should not reach higher resolution if not limited by size. The calculated Rosenthal B-factor from this dataset (49Å^2^) indicates that though the data is of high quality, an impractical number of particles would be required to achieve significantly higher resolution. For example, to improve the resolution by 0.2Å would require ∼1.6x more particles while reaching 1.2Å would require an unrealistic 71x10^6^ particles. It is possible that other benchmarks of AaLS could have improved B-factors that would match or surpass those obtained for apoferritin.

Though we did not use AFIS for this experiment, a collection speed of >300 movies per hour was achieved owing to the shorter exposure times at high magnification and the ability to use minimal beam-image shift within a hole with a very small, yet parallel beam. We anticipate that a properly calibrated AFIS routine and separation into optics groups could yield similar results. We identified a rather high magnification anisotropy (∼1.7-1.8%) at the magnification used for this dataset (as well as a similar dataset 6 months prior). Though this can be corrected for *in silico* the ultimate performance of the microscope could be compromised and requires further investigation. This highlights why benchmarking a newly installed microscope is important to any facility as no two identical hardware configurations behave exactly the same. Factory assembly, commissioning and environmental differences can affect microscope performance that can affect results at even intermediate resolution as reported before (Grant & Grigorieff, 2015).

In summary we have demonstrated that SPA at near-atomic resolution is achievable for macromolecules other than apoferritin and can be obtained from user-friendly hardware configurations located at University Core facilities. Momentum should be directed to sample optimization and novel vitrification approaches to obtain high-quality samples.

## Methods

### Protein purification

The amino-acid sequence of the Lumazine Synthase gene *from Aquifex aeolicus* (AaLS) (Uniprot O66529) fused to a Streptag at the C-terminal was codon-optimized for expression in *Escherichia coli*. The gene was ordered from Integrated DNA Technologies (IDT) and cloned using Gibson assembly. The sequence of the gene in the transformed plasmid was then verified by Sanger sequencing. The transformed BL21 (DE3) cells were grown in LB media at 37°C until reaching an optical density at 600 nm wavelength (OD_600_) of 0.8. Protein expression was induced by using 0.2 mM IPTG concentration in the culture. The expression was done at 16°C for the duration of 17 hours. The spun down cell pellet was resuspended in lysis buffer (100 mM potassium phosphate (KP_i_) buffer pH 8, 1% (v/v) Tween 20, 50 μM EDTA, and protease inhibitor). The cells were disrupted by ultrasonication and the supernatant was heated at 75°C for 45 min then spun down and filtered through 45 μm filter to remove precipitated heat-labile proteins. The filtrate was passed through StrepTrap HP 5 mL column (GE Healthcare) using 100 mM Tris-HCl pH 8.0 and 150 mM NaCl (Buffer A). AaLS was eluted by Buffer B (Buffer A + 2.5 mM d-desthiobiotin). The fractions containing the protein were collected and concentrated using 100 kDa cutoff concentrator and loaded onto Superdex 200 10/300 GL column (GE Healthcare) using 20 mM Tris-HCl pH 7.5 and 150 mM NaCl. The fractions containing the eluted protein were curated and concentrated using 100 kDa cutoff concentrator then flash frozen into liquid nitrogen and stored at -80°C.

### Electron microscopy

Frozen aliquots of AaLS were thawed on ice and diluted to 2.75 mg/ml in 20mM Tris pH 7.5, 150mM NaCl and 1mM DTT. UltrAuFoil (Quantifoil GmbH) R1.2/1.3 Au 300 mesh grids were first washed in acetone for 30 seconds followed by isopropyl alcohol for 10 seconds and left to dry. Cleaned grids were glow-discharged for 30 seconds at 30 mA in air using a PELCO easiGlow unit. A Vitrobot MKIV was then used to plunge-freeze grids in liquid ethane (3 μL sample, 2 sec blot time, Blot Force -5, 100% humidity). Grids were loaded onto a Themo Fisher Scientific Krios G4 located at the Imaging and characterization Core Lab at KAUST equipped with a cFEG, a Falcon 4i direct detection device mounted at the end of a Selectris-X post-column energy filter.

12657 movies were recorded over a 48-hour period. A nominal magnification of 270,000X resulted in a calibrated physical pixel size of 0.4553Å (calibrated as described below). The 50μm 2^nd^ condenser aperture was used and an objective aperture was omitted entirely so as not to limit the attainable resolution (100μm aperture high frequency cut-off ∼1.4Å). A dose rate of 3.1 e-/pix/sec (as measured on the detector over vacuum) resulted in a total dose of ∼46 e-/Å^2^ on the specimen over a 3 second exposure. A total of 918 frames were saved in EER format. All grid screening and data collection was carried out using EPU (version 3.5.1, Thermo-Fisher Scientific) and using stage shift rather than AFIS to minimize the effect of beam-tilt. A 5 sec stage settling time was used between stage shifts. A parallel beam diameter of 350 nm, confirmed over gold foil in diffraction mode, allowed the exposure of 9 areas within a hole using beam-image shift. A nominal defocus range of -1.2μm to -0.4μm in 0.2μm intervals was applied over the dataset and the energy filter slit width was set to 10eV without automatic re-centering of the zero-loss energy peak.

### Data processing

Prior to data collection 5 drift-correct exposures were recorded over the gold foil using the same illumination conditions as data collection. The images were then imported into magCalEM v13.0 (Dickerson *et al*., 2024). The program provides a calculated pixel size taking into account any potential magnification anisotropy of the microscope projection system. At the time of acquisition, the calibrated pixel size was measured to be 0.4553Å, 1.2% larger than the service calibrated pixel of 0.45Å.

RELION (Scheres, 2012) (version 5.0b) was used for all processing steps described below. EER format movies were gain and motion corrected using the RELION implementation with an EER fractionation group size of 18 raw frames resulting in 51 fractions with a dose per fraction of 0.88 ele/Å^2^. Following on from CTF estimation images were selected based on a resolution, defocus, figure of merit and finally manual removal of images with significant crystalline ice present. Log based autopicking was used to pick and generate a set of 2D classes for reference-based picking. In total 698,240 particles were extracted and downscaled 4x. Following on from 2D classification particles were extracted at the full pixel size in a 700-pixel box and reached a 3D auto-refine resolution of 1.97Å using icosahedral (I) symmetry. Following the 1^st^ round of CTF Refinement to estimate aberrations, magnification anisotropy and per-particle defocus the resolution improved to 1.75Å. Particle Polishing improved the resolution slightly to 1.72Å while a second round of CTF Refinement resulted in a significant increase to 1.46Å. As a final step to sort out particle heterogeneity, 3D classification without alignment and using a regularization parameter of t=10 and two classes resulted in a subset of ∼80% of particles of higher-resolution that were subjected to a final round of CTF Refinement leading to a final 3D auto-refine resolution of 1.43Å. Using the refinement run_data.star to reconstruct the particles whilst taking into consideration the Ewald Sphere resulted in a final map at 1.42Å resolution. All post-processing steps took into account the calibrated pixel size and the modulation transfer function (MTF) of the detector. The estimated beam tilt for this dataset was X=-0.02206 mrad and Y=-0.02670 mrad and an estimated magnification anisotropy of 1.75%. Subsets of particles were used for the Rosenthal B-factor estimation from an initial random selection of 240,000 from the final 470,878 particle set. Each subset was refined against a 30Å filtered reference and post-processed as above.

### Model building and refinement

Servalcat/Refmac5 (Yamashita *et al*., 2021, Murshudov *et al*., 2011) as implemented in the CCPEM suite (Burnley *et al*., 2017) was used for model refinement in combination with COOT (Emsley *et al*., 2010) for manual model building and inspection. The AaLS 1.60Å crystal structure monomer (PDB ID: 1hqk) was stripped of water molecules and docked in UCSF Chimera to one asymmetric unit of the map. Unsharpened and unweighted half maps from the Ewald Sphere corrected reconstruction and this model were used for 10 cycles of refinement with Auto symmetry set to Global and strict icosahedral (I) point group symmetry. This allowed us to work with one monomer while using a symmetry expanded model for refinement. Waters in the sharpened map were identified in COOT within a distance of 2.0-3.2Å of the protein atoms and added to this model. This hydrated model was used for another round of Servalcat refinement and the resulting difference (*F*_o_*-F*_c_) map calculated between the map and input model were masked around one asymmetric unit using RELION Mask Create and Map Process in CCPEM. PEAKMAX from the CCP4 suite (Agirre *et al*., 2023) was used to identify peaks above 2.0σ in the *F*_o_*-F*_c_ map followed by automatic selection of peaks in WATPEAK (CCP4) of less than 0.5Å from atoms in a model with added hydrogens in all possible positions. Approximately 46% of all possible hydrogens could be accounted for in the difference map (552 out of 1190).

## Acknowledgements

We would like to thanks Dr. Ashraf Al-Amoudi and Dr. Lingyun Zhao of the Imaging and Characterization Core Labs for assistance with the electron microscopes and Dr. Nagarajan Kathiresan of the Supercomputing Core lab with assistance and advise on computing. We would also like to thank Prof. Richard Henderson of MRC LMB, UK and Dr. Rafael Fernandez-Leiro of CNIO, Spain for helpful discussions. Finally, we would like to thank Joshua Dickerson and Dr. Christopher Russo of MRC LMB, UK with assistance and feedback with the magCalEM software.

## Author Contributions

The project was initiated by M.A.S and S.M.H. Lumazine synthase was purified by M.A.S. Data was collected by C.G.S and M.A.S. Data processing was done by C.G.S and M.A.S. Model refinement was done by C.G.S. The data was analyzed by C.G.S, M.A.S and A.D. C.G.S, M.A.S, A.D and S.M.H wrote the manuscript.

## Data Availability

Cryo-EM half maps have been deposited to the Electron Microscopy Data Bank (EMDB) under accession number EMD-39478. The model and raw data will be deposited to the Protein Data Bank (PDB) and Electron Microscopy Public Image Archive (EMPIAR) respectively.

## Supplementary Figures

**Supplementary Figure 1.**
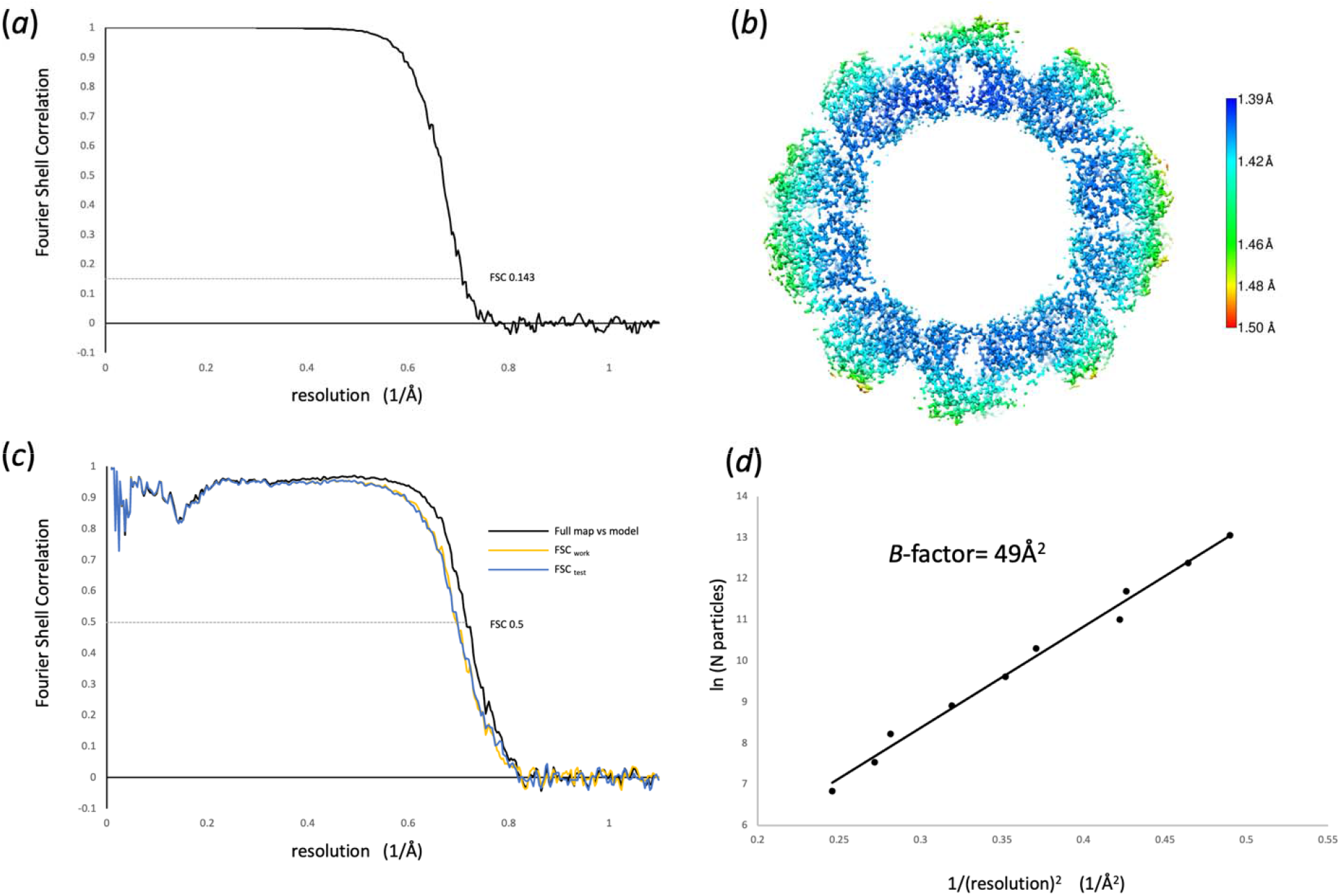
Quality analysis of the AaLS data, maps and model. (a) FSC between the final Ewald sphere corrected half maps. (b) Local resolution of the Ewald sphere correct map indicating resolution distribution with areas better than 1.42Å. (c) FSC between the full map and final model (black line). Cross validation FSC between the first half map and model refined into this map (FSCwork, yellow line) and the same model against the second half map (FSCtest, blue line). (d) Rosenthal B-Factor estimation. Subsets of particles from the final particle set were refined against a common reference and post-processed. In total 9 subsets were created and plotted along with the final particle set. The B-factor was calculated as 2 x slope of the linear fit at ∼49Å^2^.

