## Supplementary Figure 1 for "Structure of *Aquifex aeolicus* Lumazine Synthase by Cryo-Electron Microscopy to 1.42Å Resolution"

### Supplementary Material

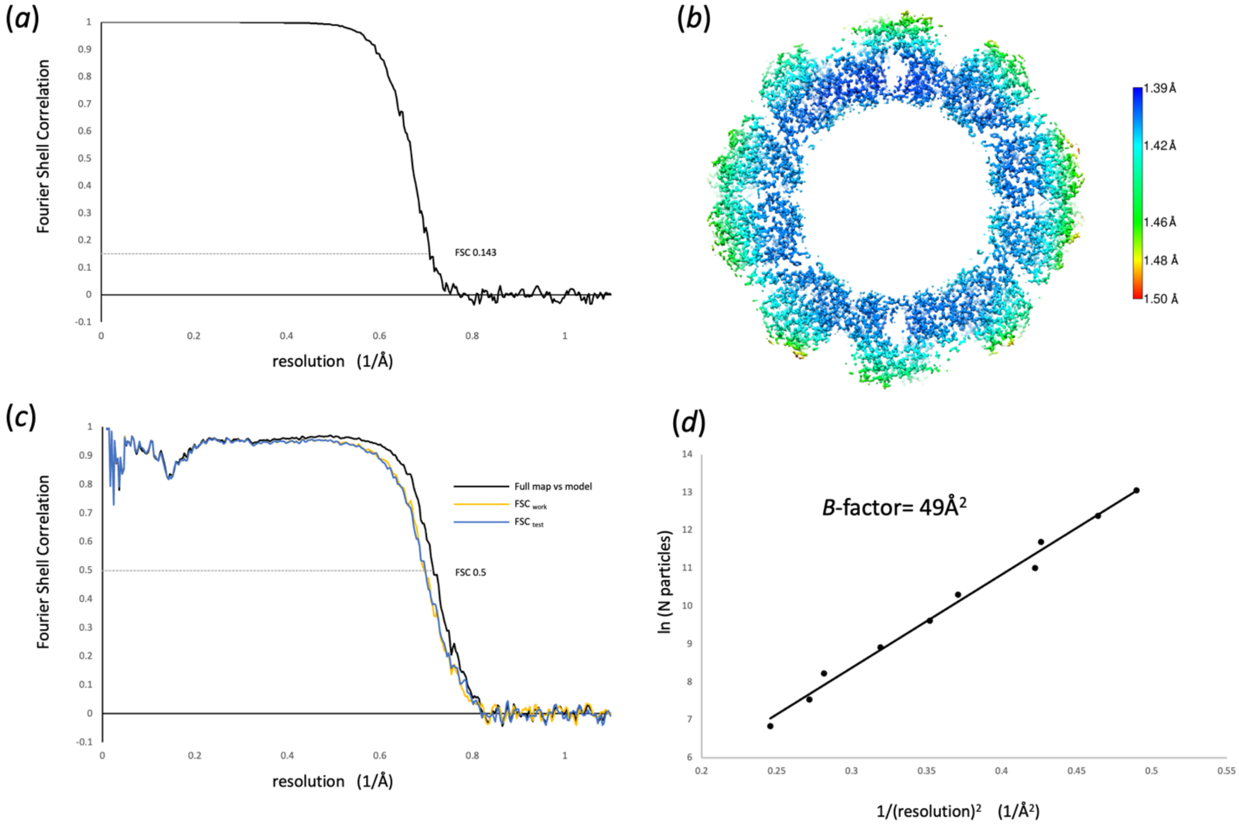


**Supplementary Figure 1.** Quality analysis of the AaLS data, maps and model. (a) FSC between the final Ewald sphere corrected half maps. (b) Local resolution of the Ewald sphere correct map indicating resolution distribution with areas better than 1.42Å. (c) FSC between the full map and final model (black line). Cross validation FSC between the first half map and model refined into this map (FSCwork, yellow line) and the same model against the second half map (FSCtest, blue line). (d) Rosenthal B-Factor estimation. Subsets of particles from the final particle set were refined against a common reference and post-processed. In total 9 subsets were created and plotted along with the final particle set. The B-factor was calculated as 2 x slope of the linear fit at ~49Å$^2$.
